# Trading accuracy for speed over the course of a decision

**DOI:** 10.1101/2020.04.02.022327

**Authors:** Gerard Derosiere, David Thura, Paul Cisek, Julie Duque

## Abstract

Humans and other animals often need to balance the desire to gather sensory information (to make the best choice) with the urgency to act, facing a speed-accuracy tradeoff (SAT). Given the ubiquity of SAT across species, extensive research has been devoted to understanding the computational mechanisms allowing its regulation at different timescales, including from one context to another, and from one decision to another. However, animals must frequently change their SAT on even shorter timescales – *i*.*e*., over the course of an ongoing decision – and little is known about the mechanisms that allow such rapid adaptations. The present study aimed at addressing this issue. Human subjects performed a decision task with changing evidence. In this task, subjects received rewards for correct answers but incurred penalties for mistakes. An increase or a decrease in penalty occurring halfway through the trial promoted rapid SAT shifts, favoring speeded decisions either in the early or in the late stage of the trial. Importantly, these shifts were associated with stage-specific adjustments in the accuracy criterion exploited for committing to a choice. Those subjects who decreased the most their accuracy criterion at a given decision stage exhibited the highest gain in speed, but also the highest cost in terms of performance accuracy at that time. Altogether, the current findings offer a unique extension of previous work, by suggesting that dynamic changes in accuracy criterion allow the regulation of the SAT within the timescale of a single decision.

**New and noteworthy:** Extensive research has been devoted to understanding the mechanisms allowing the regulation of the speed-accuracy tradeoff (SAT) from one context to another and from one decision to another. Here, we show that humans can voluntarily change their SAT on even shorter timescales – *i*.*e*., over the course of a decision. These rapid SAT shifts are associated with dynamic adjustments in the accuracy criterion exploited for committing to a choice.

## INTRODUCTION

Humans and other animals are motivated to make choices that maximize their reward rate (Simen et al. 2009; Bogacz et al. 2010a; Balci et al. 2011; Vassiliadis and Derosiere 2020). Paradoxically, while decision accuracy increases the likelihood of getting rewards, the long deliberation time necessary to make accurate choices can ultimately reduce the reward rate (Gold and Shadlen 2002; Drugowitsch et al. 2012; Carland et al. 2019). Hence, animals always need to balance the desire to gather sensory information (to make the best choice) with the pressure to act quickly, facing a speed-accuracy tradeoff (SAT; Henmon 1911; Palmer et al. 2005; Rinberg et al. 2006; Trimmer et al. 2008; Chittka et al. 2009; Salinas et al. 2014). Given the central role of the SAT in decision-making, extensive research is being devoted to understanding the computational mechanisms at the basis of its regulation (Bogacz et al. 2010b; Schall 2019, for review see Heitz 2014).

For decades, models of decision-making have offered theoretical accounts of how the brain may regulate the SAT (Stone 1960; Ratcliff and Rouder 1998; Ratcliff et al. 2001, 2003; Thapar et al. 2003; Bogacz et al. 2010a, for review see Heitz 2014). Traditional formalizations postulate that decision-making involves an accumulation of sensory evidence, which drives neural activity up to a fixed level, and once this critical threshold is reached, an action is selected (*e*.*g*., Vickers 1970; Reddi and Carpenter 2000; Usher and McClelland 2001; Leon and Shadlen 2003; Mazurek et al. 2003; Gold and Shadlen 2007; Brown and Heathcote 2008; Heitz and Schall 2012; Hanks et al. 2014; Kelly and O’Connell 2015; Derosiere et al. 2018; Alamia et al. 2019; Schall 2019). According to this simple model, to achieve a desired accuracy criterion, the brain controls the height of a neural threshold that determines how much neural activity related to evidence is needed to commit to a decision (see Mante et al. 2013; Meister et al. 2013; Marques et al. 2020, for examples of more biologically-compatible mechanisms supporting SAT regulation during decision-making). Fast decisions involve low accuracy criteria, reducing the amount of evidence required for neural activity to reach the threshold, while longer and accurate deliberations imply higher accuracy criteria. Such adaptations were shown to occur both from one SAT context to another (*e*.*g*., Forstmann et al. 2008; Herz et al. 2016, 2017) and from one decision to another within the same context (*e*.*g*., Purcell and Kiani 2016; Thura et al. 2017; Fischer et al. 2018; Desender et al. 2019), providing a key mechanism to trade speed with accuracy at different time-scales. Yet, some recent work questioned the accuracy (*e*.*g*., Ditterich, 2006) and the generalizability (*e*.*g*., Cisek et al., 2009, Thura 2016) of perfect accumulator models, as well as their suitability to describe the neural processes underlying SATs (Rae et al. 2014; Servant et al. 2019).

About a decade ago, several studies demonstrated that the amount of evidence required to commit to a choice can sometimes decrease over the course of a decision, indicative of an accuracy criterion that wanes as time elapses (*i*.*e*., rather than being fixed over time; *e*.*g*., Cisek et al. 2009; Gluth et al. 2012). To explain these data, some authors incorporated a time-dependent “urgency” signal in the decision-making models, which – combined with sensory evidence – pushes neural activity upwards over time, effectively implementing a dropping accuracy criterion (Ditterich 2006; Churchland et al. 2008; Cisek et al. 2009; Standage et al. 2011b; Drugowitsch et al. 2012; Thura et al. 2012; Kira et al. 2015). Such urgency-based decision models have been shown to better account for reward rate maximization and to better explain behavioral data than classic accumulation to static threshold models (Ditterich 2006; Churchland et al. 2008; Standage et al. 2011b; Thura et al. 2012; Drugowitsch et al. 2012; Carland et al. 2016; Murphy et al. 2016; Hauser et al. 2017; Malhotra et al. 2018; Palestro et al. 2018; Steinemann et al. 2018; although see: Hawkins et al. 2015; Voskuilen et al. 2016). If we assume the urgency signal is linear, then it can be characterized by an initial state and a growing rate, which determine the initial height and the dropping rate of the accuracy criterion, respectively, playing thus a central role in the regulation of the SAT. Consistently, in situations where speed is of essence, both the initial state (*e*.*g*., Steinemann et al., 2018; Thura, 2020; Thura et al., 2014) and the growing rate (*e*.*g*., Hanks et al., 2014; Murphy et al., 2016) of urgency are higher, implying a lower initial criterion that quickly decays over time, compared to when the emphasis is on accuracy.

Making decisions in dynamic environments sometimes requires adjusting the SAT on very short timescales – *i*.*e*., not only from one context or one decision to another but also during an ongoing decision (Gluth et al. 2012). Imagine for instance a monkey foraging for fruits in a tree, calmly evaluating which looks tastier when, all of a sudden, a more dominant monkey shows up. In such a scenario, the foraging animal will have to speed up its decision, which may lead it to commit to a choice that does not meet its initially high standards. This situation illustrates how animals sometimes need to quickly change their decision policy and expedite a decision as it unfolds. However, very little is known about the computational mechanisms that allow such rapid adjustments.

Here, we address the hypothesis that human subjects can modify their SAT at specific stages of the deliberation process, dynamically changing their accuracy criterion by either increasing or decreasing urgency. We tested this idea by assessing the behavior of 15 healthy participants in a modified version of the tokens task (Cisek et al., 2009), where penalty changes occurring halfway through the trial promoted rapid SAT shits, either in the early or in the late stage of the decision process.

## MATERIALS AND METHODS

### Participants

We tested 15 participants for this study (11 women; 24 ± 4.1 years old), recruited from the Research Participant Pool at the Institute of Neuroscience of UCLouvain. All subjects were right-handed according to the Edinburgh Questionnaire (Oldfield 1971) and had normal or corrected-to-normal vision. None of the participants had any neurological disorder or history of psychiatric illness or drug or alcohol abuse, or were on any drug treatments that could influence performance. Participants were financially compensated for their participation and earned additional money depending on their performance on the task (see below). The protocol was approved by the institutional review board of the Université catholique de Louvain, Brussels, Belgium, and required written informed consent.

### Experimental setup

Experiments were conducted in a quiet and dimly lit room. Subjects were seated at a table in front of a 21-inch cathode ray tube computer screen. The display was gamma-corrected and its refresh rate was set at 100 Hz. The computer screen was positioned at a distance of 70 cm from the subject’s eyes and was used to display stimuli during a decision-making task. Left and right forearms were placed on the surface of the table with both hands on a keyboard positioned upside-down. Left and right index fingers were located on top of the F12 and F5 keys, respectively (Figure 1.A).

**Figure 1:**
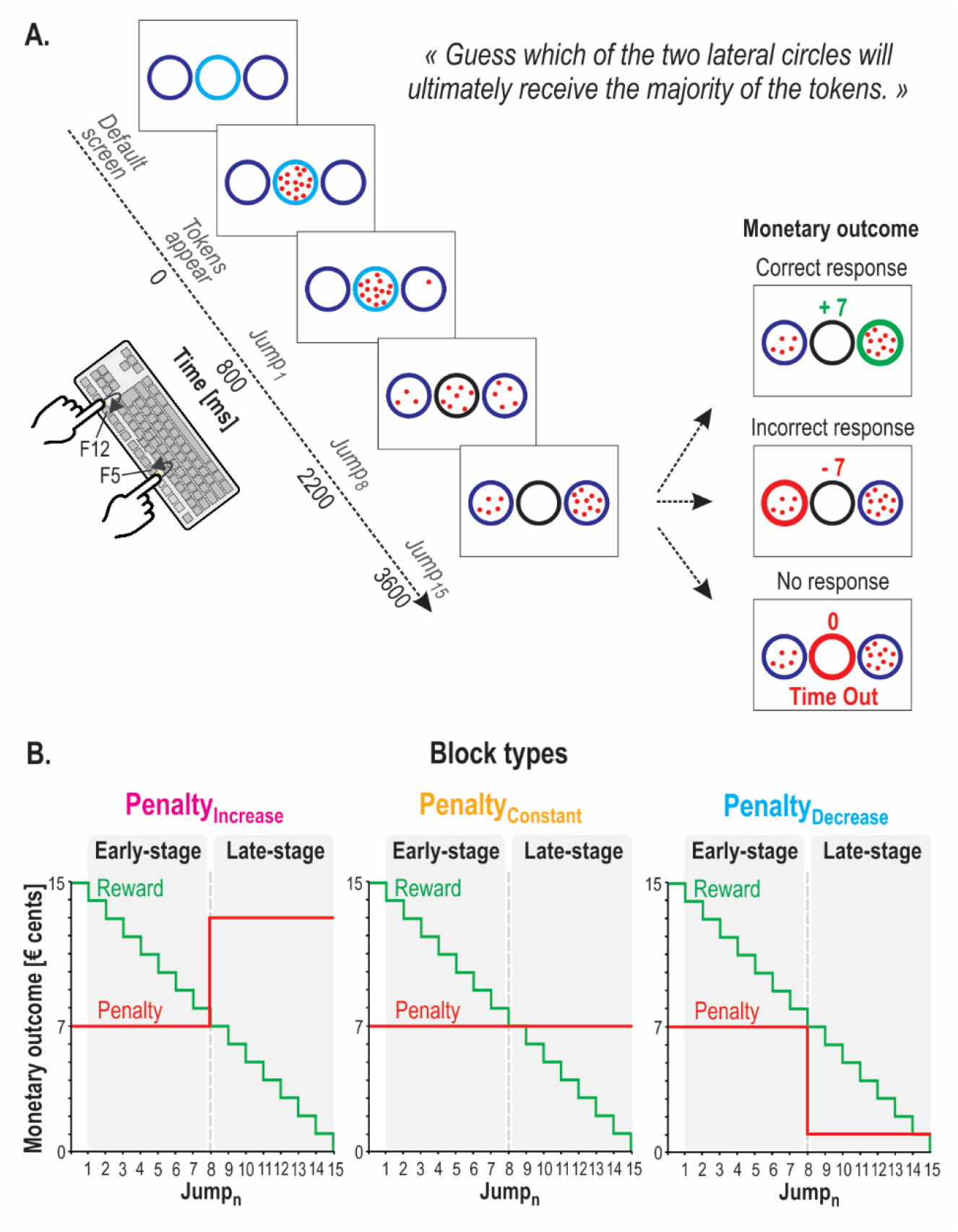
A. Schematic of the tokens task. In each trial, 15 tokens jumped one-by-one every 200 ms from the central circle to one of the lateral circles. The subjects had to indicate by a left or right index finger keypress (*i*.*e*., F12 and F5 keys, respectively) which lateral circle they thought would receive more tokens at the end of the trial. For a correct response, the subjects won, in € cents, the number of tokens remaining in the central circle at the time of the response. Hence, the reward earned for a correct response decreased over time, as depicted in B. The example presented on upper inset at the right of panel A represents a correct response provided between Jump_5_ and Jump_6_ – *i*.*e*., the score indicates that 7 tokens remained in the central circle at the moment the right circle was chosen. In contrast, as illustrated on the middle inset of A, subjects lost money if they chose the incorrect lateral circle: they received a negative score that depended on the block type, as indicated in B. In the absence of any response (“Time Out” trial, bottom inset), subjects were neither rewarded, nor penalized (score = 0). For representative purposes, the “Time Out” message is depicted below the circles in this example, while it was presented on top of the screen in the actual experiment. **B. Block types**. Incorrect responses led to a negative score, which differed in three block types. The penalty for an incorrect response always equaled 7 cents in the first half of the trial (*i*.*e*., up to Jump_8_), regardless of the block type. However, in the second half of the trial (*i*.*e*., after Jump_8_), it could then either increase to 13 cents (Penalty_Increase_ blocks; magenta, left), remain constant at 7 cents (Penalty_Constant_ blocks; yellow, center) or decrease to 1 cent (Penalty_Decrease_ blocks; blue, right). The passage from the first half of the trial (called early-stage) to the second half (late-stage) was indicated to the subjects by a change in the color of the central circle, which always turned black at Jump_8_ (see A).

### Task

The task used in the current study is a variant of the “tokens task” (Cisek et al. 2009) and was implemented by means of LabView 8.2 (National Instruments, Austin, TX). The sequence of stimuli is depicted in Figure 1.A. In between trials, subjects were always presented with a default screen consisting of three empty circles (4.5 cm diameter each), placed on a horizontal axis at a distance of 5.25 cm from each other. The central and lateral circles were light blue and dark blue, respectively, and were displayed on a white background for 2500 ms. Each trial started with the appearance of fifteen randomly arranged tokens (0.3 cm diameter) in the central circle. After a delay of 800 ms, the tokens began to jump, one-by-one every 200 ms from the center to one of the two lateral circles (*i*.*e*., 15 token jumps; Jump1 to Jump15). The subjects were instructed to indicate by a left or right index finger keypress which lateral circle they thought would ultimately receive the majority of the tokens (F12 or F5 key-presses for left or right circle, respectively). They could respond as soon as they felt sufficiently confident, as long as it was after Jump_1_ had occurred and before Jump_15_. Once a response was provided, the tokens kept jumping every 200 ms until the central circle was empty. At this time, the selected circle was highlighted either in green or in red depending on whether the response was correct or incorrect, respectively, providing the subjects with a feedback of their performance; the feedback also included a numerical score displayed above the central circle (see below, *Reward, penalty and block types* section). In the absence of any response before Jump_15_, the central circle was highlighted in red and a “Time Out” (TO) message appeared on top of the screen, together with a “0” (score, see below) above the central circle. The feedback screen lasted for 500 ms and then disappeared at the same time as the tokens did (the circles always remained on the screen), denoting the end of the trial. Each trial lasted 6600 ms.

Although to subjects the token jumps appeared completely random, the direction of each token jump was determined a priori, producing different types of trials (*e*.*g*., “easy” or “ambiguous” trials; Cisek et al. 2009). The different trial types were randomized in the full sequence of trials. The impact of these trial types on decision behavior has been studied previously (*e*.*g*., Cisek et al., 2009; Thura et al., 2014) and this issue was not investigated here as it falls beyond the scope of the present study.

One key feature of the tokens task is that it allows one to compute, in each trial, the success probability *p*_*i*_ *(t)* associated with choosing each target *i* at any time *t*. For instance, for a total of 15 tokens, if at a particular moment in time the right target contains *N*_*R*_ tokens, whereas the left contains *N*_*L*_ tokens, and there are *N*_*C*_ tokens remaining in the center, then the probability that the target on the right will ultimately be the correct one (*i*.*e*., the success probability of guessing right) is as follows:

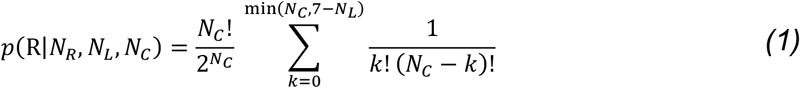

Computation of the subject’s accuracy criterion relies on the amount of sensory evidence that was available in favor of the chosen target when the subject committed to her/his choice (*i*.*e*., at decision time [DT]). Because we think it is very unlikely that subjects can calculate the real success probability function by computing a sum of factorials at every token jump (eq. 1), we computed a first-order estimation as the sum of log-likelihood ratios (SumLogLR) of individual token movements (Cisek et al., 2009). Based on the temporal profile of the accuracy criterion (*i*.*e*., of the SumLogLR at DT), it is then possible to extract an urgency function. Hence, the tokens task provides us with the possibility to estimate how the accuracy criterion, as well as urgency, vary from one experimental condition to another. Further details regarding the computation of the accuracy criterion and of the urgency function are provided later, in the *Data analyses* section.

### Reward, penalty and block types

As mentioned above, subjects received a feedback score at the end of each trial, which depended on whether they had selected the correct or the incorrect response. Correct responses led to positive scores (*i*.*e*., a reward) while incorrect responses led to negative scores (*i*.*e*., a penalty). Subjects knew that the sum of these scores would turn into a monetary reward at the end of the experiment.

In correct trials, the reward was equal to the number of tokens remaining in the central circle at the time of the response (in € cents). Hence, the potential reward for a correct response gradually decreased over time (Figure 1.B). For instance, a correct response provided between Jump_5_ and Jump_6_ led to a gain of 10 cents (10 tokens remaining in the central circle). However, it only led to a gain of 5 cents when the response was provided between Jump_10_ and Jump_11_ (5 tokens remaining in the central circle). The fact that the reward dropped over time produced an increasing urge to respond over the course of a trial, as evidenced from the urgency functions obtained in such a task (Derosiere et al. 2019).

Incorrect responses led to a negative score but here, the size of this penalty was not linearly proportional to the RT. Importantly, it differed in three block types (see Figure 1.B). The penalty for an incorrect response always equaled 7 cents in the first half of the trial (*i*.*e*., up to Jump_8_), regardless of the block type. However, in the second half of the trial (*i*.*e*., after Jump_8_), it could then either increase to 13 cents (Penalty_Increase_ blocks), remain constant at 7 cents (Penalty_Constant_ blocks) or decrease to 1 cent (Penalty_Decrease_ blocks). The passage from the first half of the trial (called early-stage) to the second half (late-stage) was indicated to the subjects by a change in the color of the central circle, which always turned black at Jump_8_. Each block type was realized on a separate experimental session (see Experimental procedure section below), and subjects were informed at the beginning of the session the block type to be realized.

We expected that the penalty shift would induce stage-specific adjustments of the SAT in the Penalty_Increase_ and Penalty_Decrease_ blocks, compared to the Penalty_Constant_ condition. Particularly, in the Penalty_Increase_ blocks, we expected that the prospect of a higher penalty at a late stage would promote faster decisions at the early stage, at the cost of accuracy. Hence, we expected subjects to trade accuracy for speed specifically when making early-stage decisions in the Penalty_Increase_ blocks. Inversely, in the Penalty_Decrease_ blocks, we predicted a tendency to make fast but less accurate decisions at a late-stage of the trial, after the drop in penalty.

### Experimental procedure

Subjects performed three experimental sessions (one for each block type) conducted on separate days at a 24-h interval. Testing always occurred at the same time of the day for a given subject, to avoid variations that could be due to changes in chronobiological states (Schmidt et al. 2006; Derosière et al. 2015). The order of the sessions was counterbalanced across participants.

The three sessions always started with two short blocks of a simple RT task. In this task, subjects were presented with the same display as in the tokens task described above. However here, instead of jumping one by one, the 15 tokens jumped simultaneously into one of the two lateral circles (always the same one in a given block) and subjects were instructed to respond as fast as possible by pressing the appropriate key (*i*.*e*., F12 and F5 for left and right circles, respectively). Because the target circle was known in advance of the block, this task did not require any choice to be made and was exploited to determine the subject’s mean “simple reaction time” (SRT) for left and right index finger responses. We obtained this SRT by computing the difference between the key-press and the time at which the 15 tokens left the central circle (Cisek et al., 2009).

Next, subjects performed training blocks to become acquainted with the tokens task. In the first one (20 trials, only run on the first session), we ran a version of the tokens task in which the feedback was simplified; the lateral circle turned either green or red, depending on whether subjects had provided a correct or incorrect response; no reward or penalty was provided here. Then, we ran two training blocks (20 trials each) in the condition subjects would be performing next during the whole session (Penalty_Increase_, Penalty_Constant_ or Penalty_Decrease_).

The actual experiment involved 8 blocks of 80 trials (640 trials per session; 1920 trials per subject). Each block lasted about 8.5 minutes and a break of 5 minutes was provided between each of them. Each session lasted approximately 120 minutes.

### Data analyses

Data were collected by means of LabView 8.2 (National Instruments, Austin, TX), stored in a database (Microsoft SQL Server 2005, Redmond, WA), and analyzed with custom Matlab (MathWorks, Natick, MA) scripts.

#### Decision time, performance accuracy, percentage of time outs

For each block type and each subject, we computed the average decision time (DT) and performance accuracy (% Correct choices over the total number of responses), as well as the percentage of “time out” trials (%TO over the total number of trials). To estimate the DT, we first calculated the reaction time (RT) during the tokens task by computing the difference between the time at which the subject pressed the key and the time of Jump_1_. We then subtracted from this RT the mean simple reaction time (SRT) obtained for each subject. This procedure allowed us to remove, from the individual RT obtained in the tokens task, the sum of the delays attributable to sensory processing of the stimulus display as well as to response initiation and muscle contraction, providing us with an estimate of the DT in the tokens task (Cisek et al. 2009; Derosiere et al. 2019).

Although the non-decision delay might differ in simple versus choice tasks (*e*.*g*., due to different encoding demands), we are always comparing DTs so those comparisons are not affected by any inaccuracies in our within-subject SRT estimate. A potential inaccuracy could occur in the calculation of the SumLogLR at DT in some trials (see below), in which we may mistakenly calculate it from the wrong token state. However, this should be quite rare given that the tokens jump only once every 200 ms, giving us a large time window.

For the analysis of DT and performance accuracy data, the dataset was split into two subsets according to whether decisions were made during the early stage (between Jump_1_ and Jump_8_ ; DTs ranging from 0 and 1400 ms) or during the late stage of the trial (between Jump_8_ and Jump_15_ ; DTs ranging from 1400 and 2800 ms). The early stage involved a total of 124 ± 15, 119 ± 16 and 150 ± 27 trials in the Penalty_Decrease_, the Penalty_Constant_ and the Penalty_Increase_ blocks, respectively, while the late stage comprised 507 ± 15, 495 ± 14 and 444 ± 25 trials in these three blocks. This split allowed us to test for the effect of the block on the subjects’ decision speed and accuracy, separately for responses provided either during the early- or during the late-stage of the trial. We predicted that, compared to the Penalty_Constant_ condition, subjects’ performance accuracy would be particularly low for responses provided during the early-stage in Penalty_Increase_ blocks and during the late-stage in Penalty_Decrease_ blocks, reflecting a propensity to trade decision accuracy for speed when the penalty is the lowest within a trial.

#### Accuracy criterion

As mentioned above, the tokens task allows us to estimate the subject’s accuracy criterion, based on the amount of evidence that was available for the chosen circle in each trial at DT. There are two possible ways to quantify this. The first is to use equation (1) to calculate the exact probability that the chosen target will be correct, given the number of tokens in each target and the number remaining in the center at the time of the choice. While this is the exact answer, we do not believe it reflects what our participants are doing because we do not believe that they can calculate equation 1, which involves a summation of factorials, every 200 ms. Thus, analyzing behavior using this quantity would presumably not reflect the processes actually used by our participants. For this reason, when analyzing behavior we quantified evidence using a different and much simpler calculation, based on the sum of the log-likelihood ratios (SumLogLR) of the individual token jumps. This is described as:

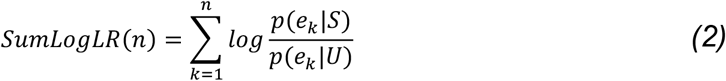

where *p(e*_*k*_ |*S)* is the likelihood of a token event e_k_ (a token jumping into either the selected or non-selected lateral circle) during trials in which the selected target is correct, and *p(e*_*k*_ |*U)* is its likelihood during trials in which the unselected target is correct. Note that p(e_k_ |S) comes out to 148620 / (16384*15) = 0.6047, while p(e_k_ |U) comes out to 97140 / (16384*15) = 0.3953, so the quantity in the summation is equal to log(0.6047/0.3953) = 0.42524 for a token that jumps to the selected target, and to the negative of that when it jumps to the other target. Consequently, equation (2) is simply a measure of the difference in the number of tokens that jumped to the selected versus unselected target, scaled by a factor of 0.42524, and does not take into account the full history of token jumps. For example, it continues to increase and decrease with each token jump even after a target has received 8 or more tokens, at which time the real probability simply saturates at 1. Also, the SumLogLR is not necessarily predictive of the correctness of the choice (*e*.*g*., in so-called “misleading trials”, subjects may respond with a positive SumLogLR after 3 token jumps but eventually end up being incorrect because the ensuing jumps eventually favor the opposite target). Nevertheless, in practice equation (2) is a very good approximation of the real probability function in most of the trials, especially for the first 10 token jumps (Cisek et al. 2009), and presumably better reflects the process used by our subjects.

To characterize the changes in accuracy criterion from one block condition to another during the early- and the late-stage of the trial, we split the SumLogLR at DT dataset into two subsets according to whether decisions were made in the former or in the latter stage. In accordance with previous studies (*e*.*g*., Cisek et al., 2009; Gluth et al. 2012; Murphy et al., 2016), we expected that the accuracy criterion would drop as the deadline to respond approached, thus leading to globally lower values in the late-relative to the early-stage of the trial. However, we predicted that, compared to the Penalty_Constant_ condition, the criterion would be particularly low during the early-stage in Penalty_Increase_ blocks and during the late-stage in Penalty_Decrease_ blocks, reflecting the subjects’ ability to adjust their criterion to a desired level at specific stages of the decision process.

#### Estimation of urgency functions

Models of decision-making incorporating an urgency signal posit that choices result from the combination of signals that reflect the available sensory evidence and the level of urgency that grows over time (*e*.*g*., Ditterich 2006; Churchland et al. 2008; Drugowitsch et al. 2012). For instance, in a minimal implementation of the urgency-gating model (Cisek et al. 2009; Thura et al., 2014; Thura 2020), noise-free evidence is multiplied by a linearly increasing urgency signal and then compared to the threshold. The result can be expressed as follows:

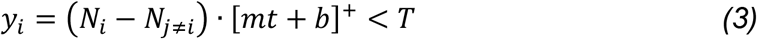

where *y*_*i*_ is the “neural activity” for choices to target *i, N*_*i*_ is the number of tokens in target *i, t* is the number of seconds elapsed since the start of the trial, *m* and *b* are the slope and y-intercept of the urgency signal, and []^*+*^ denotes half-wave rectification (which sets all negative values to zero). When *y*_*i*_ for any target crosses the neural threshold *T*, that target is chosen. Note that if we define the accuracy criterion (AC) as the amount of evidence needed to make the decision (SumLogLR at DT), then it is equal to *AC* = (*N*_*i*_(*DT*) − *N*_*j*≠*i*_(*DT*)) = *T*/[*m* · *DT* + *b*]^+^, and therefore decreases with increasing decision times.

A direct implication of such urgency-based models is that decisions made with low levels of sensory evidence (*i*.*e*., involving low accuracy criteria) should be associated with high levels of urgency and vice versa. That is, one core assumption is that a high urgency should push one to commit to a choice even if evidence for that choice is weak, effectively implementing a low accuracy criterion. Hence, the accuracy criterion values (SumLogLR at DT) can be exploited to estimate the level of urgency at DT (*e*.*g*., Thura et al., 2014; Thura, 2020).

Here, we estimated the level of urgency based on the accuracy criterion values obtained for different DTs. We first grouped the trials in bins as a function of the DT and calculated the average accuracy criterion for each bin. Nine bins were defined, with the first bin including decisions made between 800 and 1000 ms, the second bin including decisions made between 1000 and 1200 ms, and so on, until the last bin covering the period between 2400 and 2600 ms. The accuracy criterion values preceding 800 ms or following 2600 ms were not considered for this analysis because many subjects did not respond at these times (59.5 ± 0.04 % of the bins were missing values for these very early and very late times). Considering a model in which evidence is multiplied by an urgency signal, we estimated urgency values based on the accuracy criterion obtained at each bin, in each subject and each block condition, as follows:

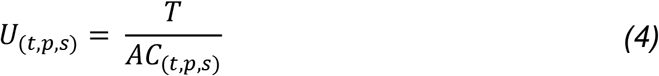

Above, *t* is the DT bin, *p* is the penalty condition, *s* is the subject number, *AC* is the accuracy criterion value (*i*.*e*., SumLogLR at DT), *T* is a constant representing a fixed neural threshold (which we fixed to 1), and *U* is the estimated urgency value. We then fitted regression models over the obtained urgency values. A linear model and a second-order polynomial model were fitted and the Akaike Information Criterion (AIC) was obtained for each subject and each block condition, allowing us to compare the two models to each other. We predicted that, compared to the Penalty_Constant_ condition, urgency would be particularly high during the early-stage in Penalty_Increase_ blocks and during the late-stage in Penalty_Decrease_ blocks, and that the polynomial model would thus better capture the dynamic changes in urgency in these two block conditions compared to the linear one (*i*.*e*., lower AIC values for polynomial fits).

### Statistical analyses

Statistica software was used for all analyses (version 7.0, Statsoft, Oklahoma, United-States). The DT, performance accuracy and SumLogLR at DT data were analyzed using two-way repeated-measure ANOVAs (ANOVA_RM_) with BLOCK (Penalty_Increase_, Penalty_Constant_, Penalty_Decrease_) and STAGE (early-stage, late-stage) as within-subject factors. The %TO data were analyzed using a one-way ANOVA_RM_ with BLOCK (Penalty_Increase_, Penalty_Constant_, Penalty_Decrease_) as a within-subject factor. When appropriate, LSD post-hoc tests were used to detect paired differences. Finally, the AIC values obtained from the urgency fits were analyzed using a two-way ANOVA_RM_ with BLOCK (Penalty_Increase_, Penalty_Constant_, Penalty_Decrease_) and MODEL (linear, polynomial) as a within-subject factors. When performing ANOVA_RM_ ’s, Maunchley’s tests were exploited systematically to check for data sphericity and Greenhouse-Geisser (GG) corrections were used to correct for any significant deviation from sphericity. Results are presented as mean ± SE.

## RESULTS

### Decision time, performance accuracy and %TO

The average DT was not significantly different in the Penalty_Increase_, the Penalty_Constant_ and the Penalty_Decrease_ blocks (1503 ± 32, 1527 ± 25 ms and 1505 ± 19 respectively), as indicated by the absence of main effect of BLOCK (F_1,14_ = 0.73, p = .489). Importantly though, the ANOVA_RM_ revealed a significant BLOCK*STAGE interaction on the DT data (F_1,14_ = 10.04, p = .0005; see Figure 2.A-B). In fact, the DT of responses provided during the early stage was significantly lower in the Penalty_Increase_ (990 ± 43 ms) than in both the Penalty_Constant_ (1061 ± 37 ms; p = .004) and the Penalty_Decrease_ blocks (1063 ± 32 ms; p = .003). Conversely, the DT of responses provided during the late stage was significantly lower in the Penalty_Decrease_ (1946 ± 15 ms) than in both the Penalty_Constant_ (1994 ± 24 ms; p = .046) and the Penalty_Increase_ blocks (2016 ± 30 ms; p = .005). Importantly though, DTs were similar for the Penalty_Increase_ and the Penalty_Constant_ blocks during the late stage (p = .353); DTs were also comparable for the Penalty_Decrease_ and the Penalty_Constant_ blocks during the early-stage (p = .925). These findings indicate that subjects increased their decision speed at very specific stages during the trial in the Penalty_Increase_ and Penalty_Decrease_ blocks: they made faster decisions specifically during the early-stage of the Penalty_Increase_ blocks and during the late-stage of the Penalty_Decrease_ blocks compared to the Penalty_Constant_ block type.

**Figure 2:**
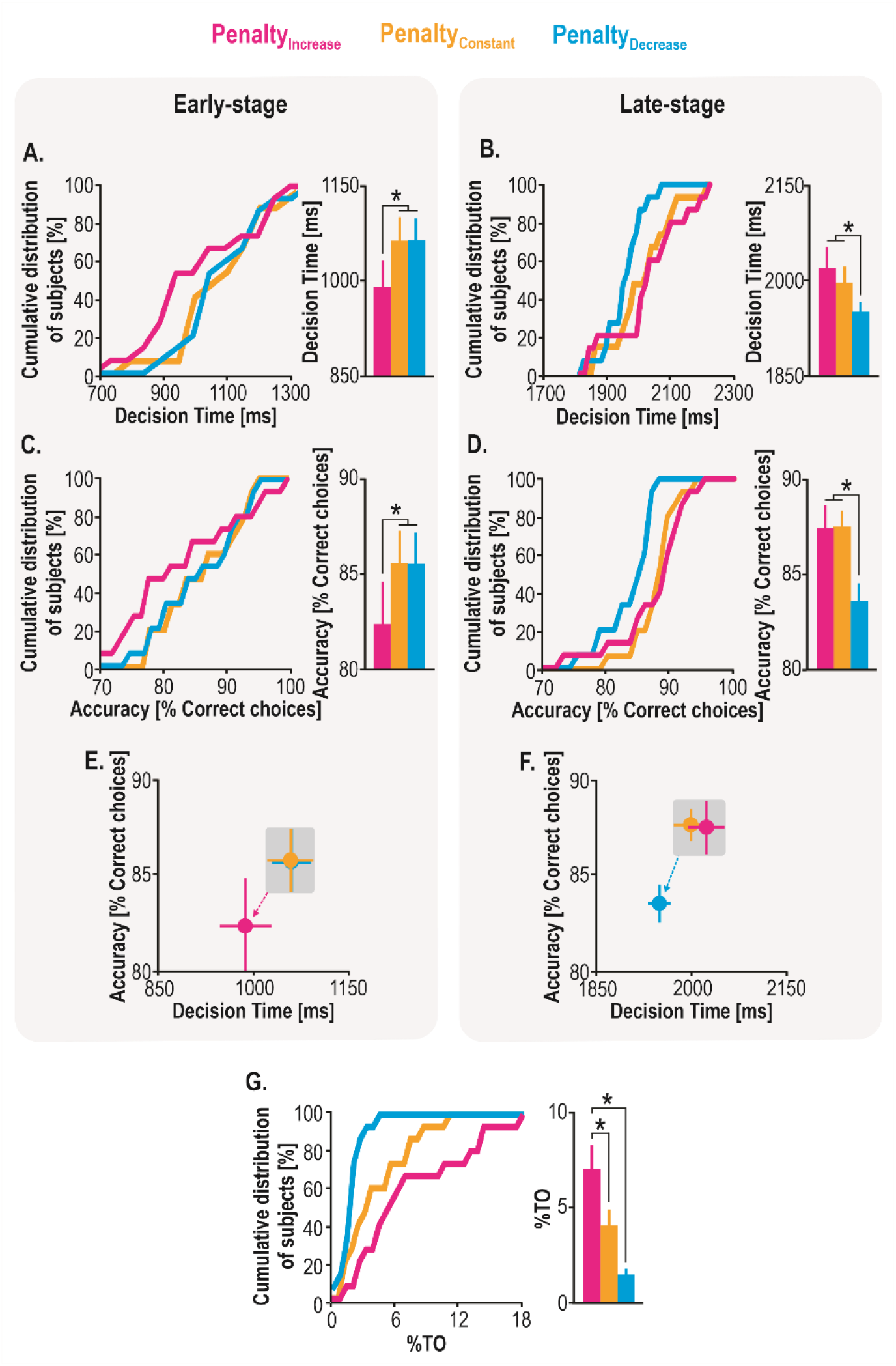
A. Decision Time (DT). Cumulative distribution of subjects and mean DT measured at the early stage of the trial in the Penalty_Increase_ (magenta traces), the Penalty_Constant_ (yellow traces) and the Penalty_Decrease_ blocks (blue traces). **B**. Same as *A*. for the late stage. **C. Accuracy (% of correct choices)**. Cumulative distribution of subjects and mean accuracy measured at the early stage of the trial in the Penalty_Increase_ (magenta traces), the Penalty_Constant_ (yellow traces) and the Penalty_Decrease_ blocks (blue traces). **D**. Same as *C*. for the late stage. **E and F. Shift in DT (x-axis) and accuracy (y-axis)**. The graphs summarize the effects represented in A and C, and B and D, respectively. A decrease in DT and in accuracy is observed specifically in the early stage of the Penalty_Increase_ block (magenta dot) and in the late stage of the Penalty_Decrease_ block (blue dot). **G. Percentage of time out trials (%TO)**. Cumulative distribution of subjects and mean %TO in the Penalty_Increase_ (magenta traces), the Penalty_Constant_ (yellow traces) and the Penalty_Decrease_ (blue traces) blocks. * Between-block difference at p < .05. Error bars represent SE.

The average performance accuracy was not significantly different in the Penalty_Increase_, the Penalty_Constant_ and the Penalty_Decrease_ blocks (84.9 ± 1.8, 86.6 ± 1.0 % and 84.6 ± 1.2, respectively; no main effect of BLOCK: F_1,14_ = 1.48, p = .244), but here again, the BLOCK*STAGE interaction was significant (F_1,14_ = 5.83, p = .008; see Figure 2.C-D). As such, responses provided during the early stage were associated with a lower accuracy in the Penalty_Increase_ (82.4 ± 2.5 %) than in both the Penalty_Constant_ (85.7 ± 1.7 %; p = .027) and the Penalty_Decrease_ blocks (85.5 ± 1.8 %; p = .038). Furthermore, responses provided during the late stage were associated with a lower accuracy in Penalty_Decrease_ (83.6 ± 1 %) than in both the Penalty_Constant_ (87.6 ± 0.9 %; p = .009) and the Penalty_Increase_ blocks (87.4 ± 1.4 %; p =.013). Importantly though, accuracy was similar for the Penalty_Increase_ and the Penalty_Constant_ blocks during the late stage of trials (p = .878), while it was comparable for the Penalty_Decrease_ and the Penalty_Constant_ blocks during the early stage (p = .879). Hence, consistent with the DT findings, subjects decreased their decision accuracy at specific stages of the trial in the Penalty_Increase_ and Penalty_Decrease_ blocks: compared to Penalty_Constant_ blocks, they made more errors during the early stage in the Penalty_Increase_ blocks and during the late stage in the Penalty_Decrease_ blocks.

Finally, the ANOVA_RM_ revealed a significant effect of BLOCK on the time out (%TO) data (GG-Corrected F_1.21,16.97_ = 15.95, p < .0001; see Figure 2.G). The %TO was indeed higher in the Penalty_Increase_ (7.2 ± 1.3 %) than in both the Penalty_Constant_ (4.1 ± 0.8 %; p = .004) and the Penalty_Decrease_ blocks (1.5 ± 0.2 %; p < .0001). In addition, it was lower in the Penalty_Decrease_ than in the Penalty_Constant_ blocks (p = .016). Hence, the lower the penalty was during the late-stage of the trial, the less the subjects were inclined to be cautious and to avoid responding, consistent with the reduced decision accuracy observed in the late-stage of Penalty_Decrease_ blocks.

In summary, these behavioral observations indicate that subjects traded decision accuracy for speed specifically at times where the penalty was lowest relative to the rest of the trial (see Figure 3 for examples of individual data). In Penalty_Increase_ blocks, the perspective of a rise in penalty promoted faster but less accurate decisions specifically in the first half of the trial, while in Penalty_Decrease_ blocks, the penalty drop promoted faster but less accurate decisions in the second half of the trial.

**Figure 3:**
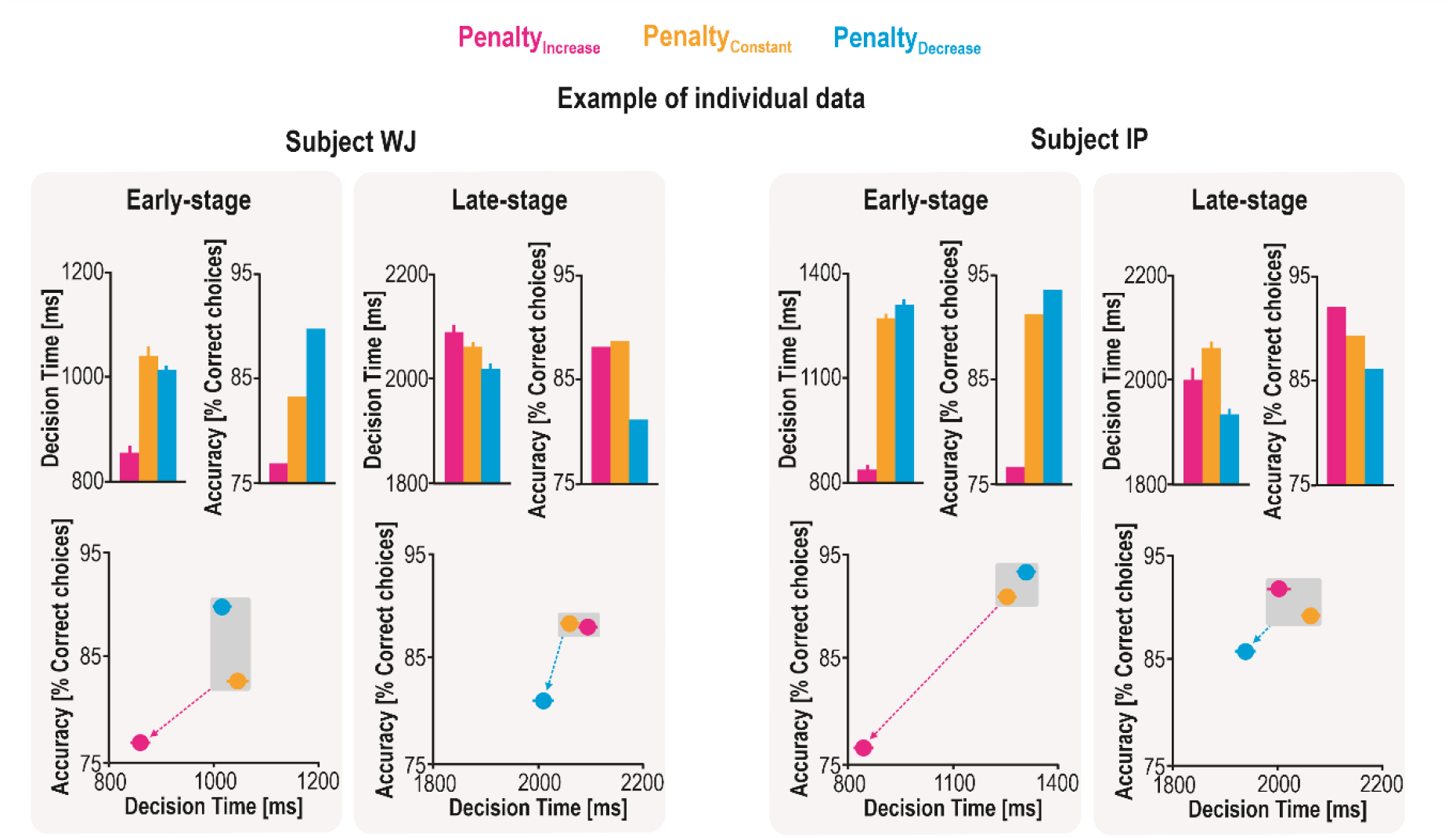
Example of individual data. DT and accuracy data are represented for the three block conditions and each stage. Error bars on DT data represent between-trial variability. Overall, these graphs highlight that the stage-specific shifts in decision speed and accuracy were evident at the single-subject level.

### Accuracy criterion and urgency functions

The accuracy criterion based on which subjects made their decision was estimated using the SumLogLR at DT: the higher the SumLogLR at DT, the higher the accuracy criterion. Overall, decisions made during the early stage were based on a higher accuracy criterion than those made during the late stage of trials (1.46 ± 0.08 and 1.18 ± 0.04 a.u., respectively), as confirmed by the ANOVA_RM_ showing a main effect of the factor STAGE on the SumLogLR at DT (F_1,14_ = 205.85, p < .0001). Hence, subjects’ accuracy criterion decreased over the course of the decision process, putatively indicating an increasing urge to respond as the central circle was emptying.

Interestingly, the accuracy criterion also depended on the BLOCK under consideration, as revealed by a significant BLOCK*STAGE interaction (F_2,28_ = 3.79, p = .035). For early-stage decisions, it was lower in the Penalty_Increase_ (1.39 ± 0.13 a.u.) than in the Penalty_Constant_ blocks (1.52 ± 0.08 a.u.; p = .026), with a drop of 10.34 ± 6.98 % (Figure 4.A). For late-stage decisions, the criterion tended to be lower in the Penalty_Decrease_ (0.62 ± 0.02 a.u.) than in the Penalty_Constant_ blocks (0.71 ± 0.03 a.u.; p = .122), with a significant drop of 10.41 ± 3.5 % (*i*.*e*., t-test against 0: t_14_ = 2.97, p = .010, percentage data were normally distributed; Figure 4.B). Importantly, these effects were stage-specific: the criterion was comparable in the Penalty_Decrease_ (1.46 ± 0.13 a.u.) and Penalty_Constant_ blocks for early-stage decisions (1.52 ± 0.08 a.u.; p = .259), as well as in the Penalty_Increase_ (0.75 ± 0.04 a.u.) and Penalty_Constant_ blocks for late-stage decisions (0.71 ± 0.03 a.u.; p = .442). Altogether, these findings indicate that subjects were able to lower their criterion for committing to a choice at specific stages of the trial, in a dynamic way.

**Figure 4:**
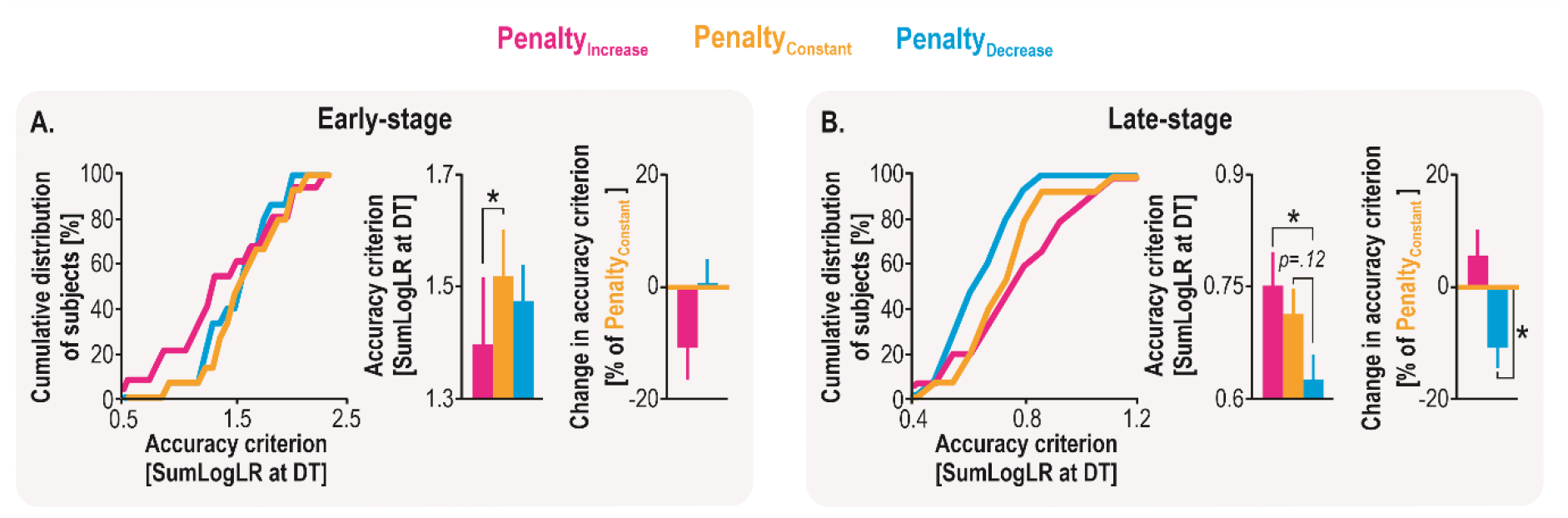
Accuracy criterion (*i.e*., the SumLogLR at DT). **A**. Cumulative distribution of subjects and mean accuracy criterion values measured at the early stage of the trial in the Penalty_Increase_ (magenta traces), the Penalty_Constant_ (yellow traces) and the Penalty_Decrease_ blocks (blue traces). **B**. Same as *A*. for the late stage. * : Between-block significant difference at p < .05. Error bars represent SE.

These stage-specific adjustments in accuracy criterion appeared to have a significant impact on decision speed and accuracy. Indeed, we observed significant positive correlations between the block-related adjustments in criterion and the block-related shift in DT as well as in performance accuracy, both when considering the early-stage (*i*.*e*., [100-(Penalty_Increase_ /Penalty_Constant_*100)] and the late-stage decisions [100-(Penalty_Decrease_ /Penalty_Constant_*100)]; all R-values = [.56 .84], all p-values = [.0312 .00008]; Figure 5). That is, the subjects who decreased the most their criterion in one block type with respect to the Penalty_Constant_ condition were those who presented the highest gains in decision speed, but also the highest costs in terms of accuracy.

**Figure 5:**
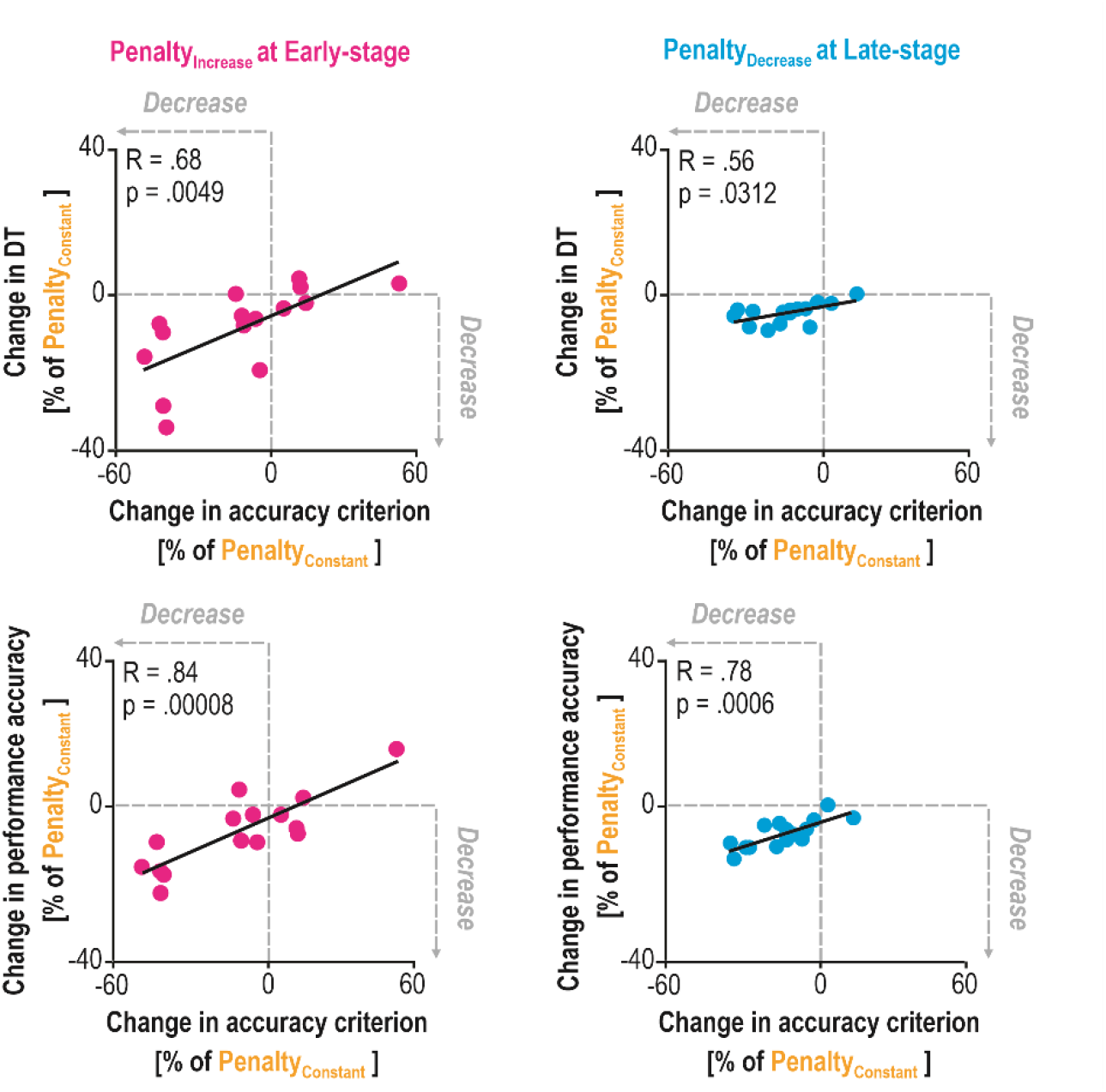
Relationships between the shift in accuracy criterion (x-axis) and the shift in DT and accuracy (y-axes; top and bottom, respectively). A significant positive correlation was found between the block-related adjustments in criterion (*i*.*e*., [100-(Criterion_PenaltyIncrease_ /Criterion_PenaltyConstant_*100)] for early-stage decisions and [100-(Criterion_PenaltyDecrease_ /Criterion_PenaltyConstant_*100)] for late-stage decisions) and the block-related shift in DT and accuracy.

A direct implication of urgency-based models is that decisions made with a low accuracy criterion are associated with a high level of urgency and vice versa. Hence, we used the temporal profile of the accuracy criterion, obtained for decisions made between 800 and 2600 ms (presented in Figure 6.A), to estimate urgency functions. Linear and polynomial models were fitted over the rectified SumLogLR at DT values and AIC were obtained for each model (Figure 6.B).

**Figure 6.**
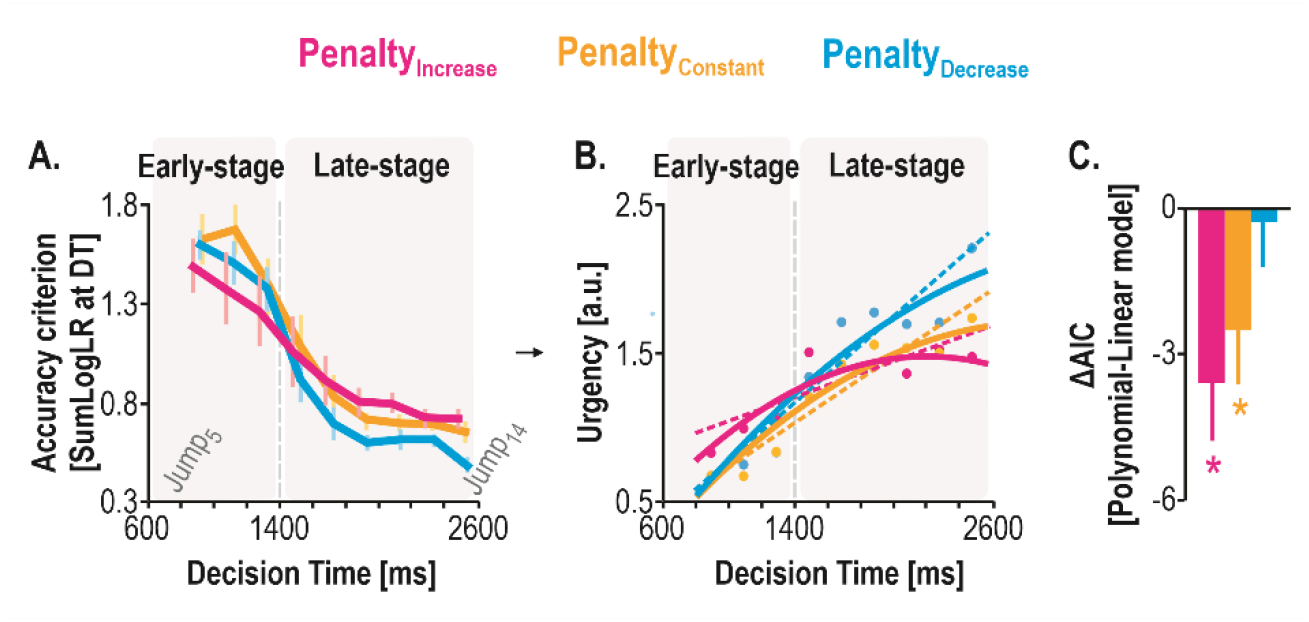
Urgency estimation. A. Temporal evolution of the accuracy criterion as a function of DT. B. Urgency functions. Functions were obtained using a linear and a second-order polynomial model fit of the rectified SumLogLR at DT data presented in A. Overall, this analysis showed that urgency increased as time elapsed over the course of a trial. Most interestingly, urgency was higher in the Penalty_Increase_ than in Penalty_Constant_ block in the early stage while it was lower in the former than in the latter block condition in the late stage. Further, while urgency was comparable in the Penalty_Decrease_ and in the Penalty_Constant_ blocks in the early stage, it became higher in the former than in the latter block in the late stage, showing thus a stage-dependent adjustment. **C. Delta AIC values**. AIC values obtained for the polynomial model were subtracted from values obtained for the linear model. The graph shows that AIC values were significantly lower for polynomial than for linear models in the Penalty_Constant_ and Penalty_Increase_ blocks, but not in the Penalty_Decrease_ block, probably due to the tendency of urgency to drop as the deadline to respond drew nearer the two former block conditions (see B). * : Between-model significant difference at p < .05. Error bars represent SE.

As evident in Figure 6.B, this analysis showed that urgency increased as time elapsed over the course of a trial, consistent with previous observations (*e*.*g*., Thura et al. 2014; Murphy et al. 2016; Hauser et al. 2018; Derosiere et al. 2019). Most interestingly, urgency was higher in the Penalty_Increase_ than in Penalty_Constant_ block in the early stage while it was lower in the former than in the latter block condition in the late stage. Besides, while urgency was comparable in the Penalty_Decrease_ and in the Penalty_Constant_ blocks in the early stage, it became higher in the former than in the latter block in the late stage, thus showing a stage-dependent adjustment.

Further, the ANOVA_RM_ performed on the AIC values revealed a BLOCK*MODEL interaction (F_2, 28_ = 3.53, p = .043. Figure 6.C). The AIC values were lower for the polynomial than for the linear model in the Penalty_Constant_ (0.18 ± 2.11 and 2.75 ± 2.34, respectively; p = .007) and Penalty_Increase_ blocks (0.49 ± 3.59 and 4.11 ± 3.21, respectively; p = .0003), but not in the Penalty_Decrease_ block (6.99 ± 2.11 and 7.34 ± 2.39, respectively; p = .700). Hence, the polynomial model captured more variance of the changes in urgency specifically when considering the Penalty_Constant_ and the Penalty_Increase_ data. As such, urgency tended to drop as the deadline to respond drew nearer in the late stage of these two block conditions.

## DISCUSSION

In dynamic environments, humans and other animals often need to change their choice SAT while a decision is ongoing. However, very little is known about the computational mechanisms that allow these rapid changes of decision policy. In the present study, we addressed the hypothesis that human subjects can shift their SAT at specific stages of the deliberation process, by dynamically adjusting their accuracy criterion. Participants performed a modified version of the tokens task (Cisek et al., 2009), where an increase or a decrease in penalty occurring halfway through the trial promoted rapid SAT shifts, either in the early or in the late decision stage. Our results reveal that subjects traded accuracy for speed specifically at times where the penalty was the lowest within a trial. Interestingly, these changes were accompanied by stage-specific adjustments in their accuracy criterion; in fact, those who decreased the most their criterion presented the highest gains in decision speed, but also the highest costs in terms of accuracy.

Several studies have now revealed the flexibility with which humans can adapt their choice SAT at different time-scales, including from one context to another (*e*.*g*., Palmer et al. 2005; Forstmann et al. 2008; Ratcliff and McKoon 2008; Herz et al. 2016, 2017) and from one decision to another (*e*.*g*., Purcell and Kiani 2016; Thura et al. 2017; Fischer et al. 2018; Desender et al. 2019). The current findings offer a unique extension of this work, by showing that the SAT can be modulated on an even shorter time-scale – *i*.*e*., over the course of a single decision. In Penalty_Increase_ blocks, decisions were faster but less accurate in the first half of the trial (*i*.*e*., compared to the Penalty_Constant_ condition), while in Penalty_Decrease_ blocks, such SAT shifts occurred in the second half of the trial. The occurrence of a shift in the first half of the trial in Penalty_Increase_ blocks indicates the operation of a proactive, anticipatory process, through which the prospect of a future rise in penalty determined the decision policy to adopt for early-stage decisions. As such, in the current task, subjects likely chose a policy for modifying their SAT before the trial had even started (or before the block of trials). Given that each block (and even each session) always involved the same type of penalty change, subjects could determine what decision policy they should adopt in this specific setting and apply it during deliberation. Whether rapid shifts in SAT can occur reactively (*e*.*g*., following online, unpredictable cues) remains an open question, worthy of future investigation.

Individuals’ accuracy criterion dropped over time in all block conditions, consistent with the idea of an urgency signal pushing subjects towards commitment as time elapses (Ditterich 2006; Churchland et al. 2008; Standage et al. 2011; Murphy et al. 2016; Malhotra et al. 2017; Steinemann et al. 2018; Hauser et al. 2018; Palestro et al. 2018; Spieser et al. 2018; Derosiere et al. 2019; Thura 2020; Shinn et al. 2020). Moreover, the temporal dynamics of this drop depended on whether the penalty increased or decreased halfway through the decision process. In the Penalty_Increase_ blocks, subjects lowered their accuracy criterion specifically in the early decision stage (*i*.*e*., relative to Penalty_Constant_ blocks) while in Penalty_Decrease_ blocks, they did so in the late decision stage. In fact, the adjustment of the accuracy criterion was more pronounced in the early decision stage (*i*.*e*., in the Penalty_Increase_ relative to the Penalty_Constant_ blocks), than in the late one (*i*.*e*., in the Penalty_Decrease_ relative to the Penalty_Constant_ blocks, where differences were only marginally significant). Hence, participants seemed more effective at adjusting their accuracy criterion for early-compared to late-stage decisions. One possible explanation for this is that the accuracy criterion was inherently higher for early decisions than for late ones, leaving more room for volitional regulation. Alternatively, it may be the case that the incentive to adjust the accuracy criterion was stronger in the early stage of Penalty_Increase_ blocks than in the late stage of Penalty_Decrease_ ones. As such, because of the natural aversion of humans to risk (Weber et al. 2004; Zhang et al. 2014), the prospect of a future rise in penalty might have been more salient than the sudden drop in penalty, thus leading to stronger changes in the accuracy criterion in the former block condition.

Our model comparison also supports the idea that subjects dynamically adjusted their level of urgency over time. Indeed, the polynomial model captured more variance of the changes in urgency when considering the Penalty_Constant_ and the Penalty_Increase_ data (*i*.*e*., compared to a linear model), but not when considering the Penalty_Decrease_ data. As such, urgency tended to drop as the deadline to respond drew nearer in the Penalty_Constant_ and the Penalty_Increase_ blocks. This effect is putatively due to the fact that the net value of responding became lower than the value of not responding as the deadline approached in these two block conditions, but not in the Penalty_Decrease_ condition, encouraging subjects to lower their level of urgency dynamically in order to refrain from responding. This interpretation is supported by the observation that subject presented a higher percentage of time out trials in the Penalty_Constant_ and the Penalty_Increase_ blocks relative to the Penalty_Decrease_ one.

Importantly, the shape of urgency signals is likely dependent on multiple contextual factors (such as risk here), on individual traits (Carland et al., 2019) and on the type of task and behavior at play. While our findings suggest that urgency signals can take the form of non-linear ramps in decision-making tasks involving dynamic SAT adjustments, such signals might include even more intricate shapes such as pulses or pauses to construct complex motor behaviors in other task settings, such as when moving to a beat.

Overall, the present study builds on former work on the computational mechanisms underlying the SAT policy. Consistent with past research, we show that the accuracy criterion progressively drops over time during the decision process, in line with an increased urge to commit as the time left to respond diminishes. Most importantly, we provide evidence that rapid shifts in SAT can occur over the course of an ongoing decision and that these changes could be related to dynamic adjustments of the accuracy criterion (Shinn et al., 2020). Future work is needed to extend the current observations to situations involving reactive SAT shifts, which may emerge in response to online sensory cues.

## SOURCE DATA

All raw data and processed data exploited in this study are available at: https://osf.io/wj4z7/

## Acknowledgements

This work was supported by grants from the “Fonds Spéciaux de Recherche” (FSR) of the Université Catholique de Louvain, the Belgian National Funds for Scientific Research (FRS-FNRS: MIS F.4512.14). GD was a postdoctoral fellow supported by the FNRS (FNRS 1B134.18). We thank Simon Van Hemelrijck and Julien Grandjean for their help in the acquisition of the data.

